# Background environment modulates motor contagions in humans

**DOI:** 10.1101/2023.03.31.535099

**Authors:** Hiroto Saito, Kentaro Fukuchi, Masahiko Inami, Gowrishankar Ganesh

**Affiliations:** Research Center for Advanced Science and Technology, The University of Tokyo, 4-6-1 Komaba, Meguro-ku, Tokyo, 153-8904, Japan; School of Interdisciplinary Mathematical Science, Meiji University, 4-21-1 Nakano, Nakano-ku, Tokyo 164-8525 Japan; UM-CNRS Laboratoire d’Informatique de Robotique et de Microelectronique de Montpellier (LIRMM), 161, Rue Ada, Montpellier, France

## Abstract

Motor contagions refer to implicit effects in one’s actions induced by the observation of actions made by others. A plethora of studies over the last two decades have exhibited that observed, as well as predicted, actions can induce various kinds of motor contagions in a human observer. However, motor contagion has always been investigated in regard to different features of an observed action and it remains unclear whether the environment, in which an observed action takes place, modulates motor contagions as well. Here we investigated the effect of the observed environment on motor contagions using an empirical hand steering task in which the participants were required to move a cursor through visual channels of different shapes. We observed the movement time of observers to be influenced by both the movement of the cursor they observed, as well as the background (channel shape) in which the cursor movement was observed. Observers consistently made faster movements after observing steering movements in a ‘narrowing’ channel compared to a ‘widening’ channel. These results show a distinct effect of the environment, in which an observed action occurs, on one’s own movement.

## Introduction

Human motor behavior is implicitly influenced by the observation of actions of others, effects referred to as *motor contagions*^1, 2^. Motor contagions exhibit how our behaviors can be influenced by others near us and provide an interesting insight into how the visual and motor systems interact in our brains. These effects have thus attracted a lot of attention in sports science^3–8^ as well as neuroscience^1, 9–11^.

Most of the motor contagions investigated in the literature have been so-called *action-imitative contagions* (AICs)^12^ which are induced simply by the observation of actions and have a signature characteristic—they cause certain features of one’s action to become similar to that of the observed action. These features include in terms of kinematics^1, 13, 14^, outcome^3–5^, goal or intention^15^, or overall behavior^16^. On the other hand, recent studies have exhibited the presence of a second type of motor contagion, namely, *prediction error–induced contagions* (PECs), which are not just determined by the actions observed, but also by what the observer predicted the actions to be^7, 8, 12, 17^. However, all previous motor contagion studies have investigated the effect of observed actions on one’s behavior, and it remains unclear whether and how the observed environment, in which an action takes place, effects one’s behavior as well. In particular, it can be argued that the prediction of actions requires one to consider the environment and hence that PECs exhibit an effect of the environment, but previous PEC studies have only studied action modulations within one static environment, and have not considered modulations in the environment itself.

Here, to investigate the effect of the background environment on motor contagions, we evaluated an empirical hand *steering task*^18, 19^ where participants were required to move a cursor through a visual channel of constant width (C_const_). We collected the cursor movement made by actors in two channels that were of a different shape: C_wide_ channels, which became wider along the direction of the movement, and C_nar_ channels, which became narrower along the direction of the movement. The C_wide_ channel is characterized by human movements that tend to accelerate across time (M_wide_) while the C_nar_ is characterized by movements that tend to deaccelerate across the length of the channel (M_nar_)(Figure 2, 3). The participants were then shown scenarios in a 2 × 2 design in which the two channels (C_wide_ and C_nar_) were mixed with the two movements (M_wide_ and M_nar_) made by the actors, and we evaluated how observing the movements in the two environments (channels) effect the contagions induced in the participant’s own cursor movements in the C_const_ channels.

**Figure 1.**
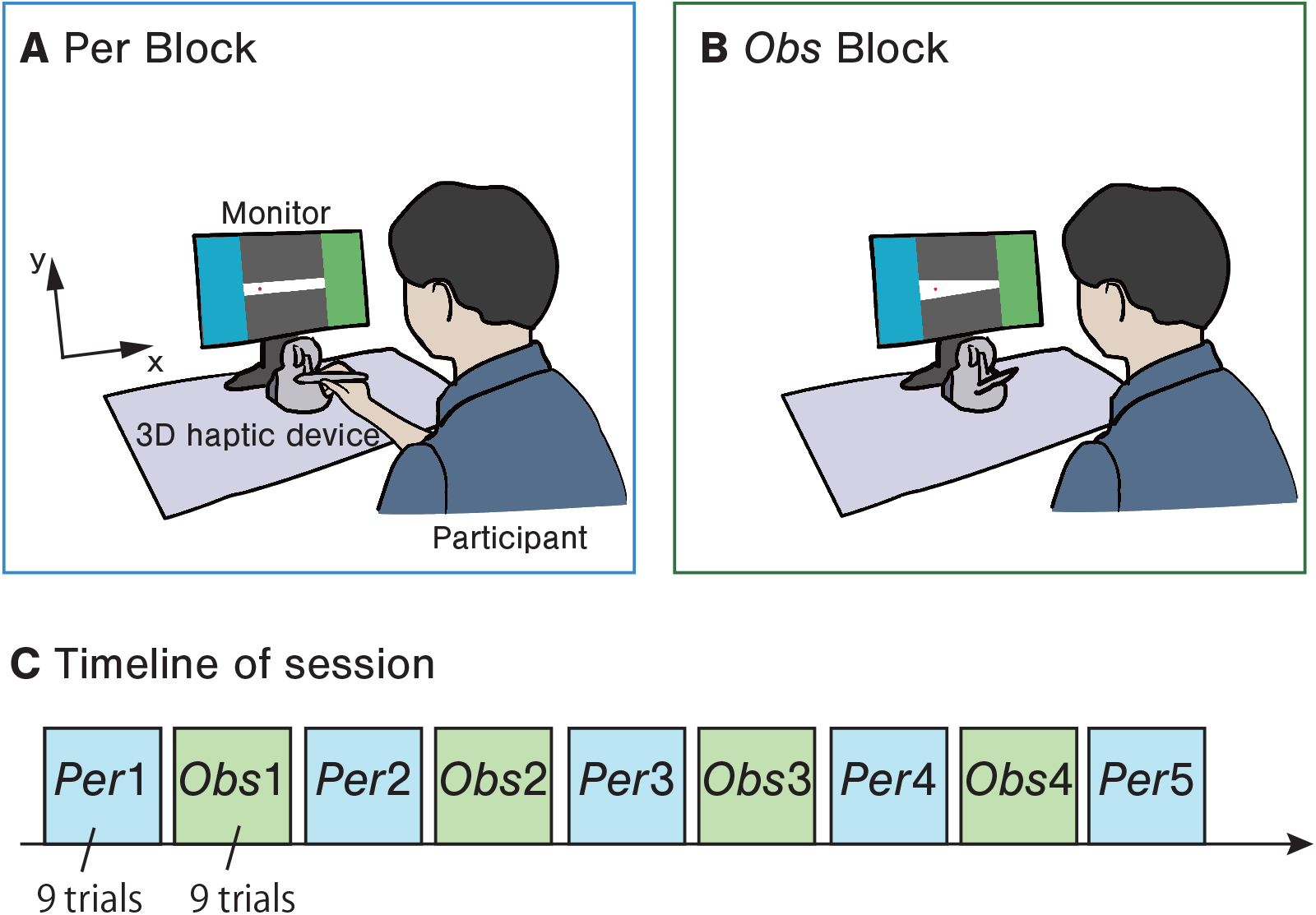
The experiments consist of two types of blocks. (A) In the *Per* block, the participants performed the steering task with a straight channel as quickly as possible. (B) In the *Obs* block, the participants watched videos of the cursor motion by unknown actors who performed steering tasks with the narrowing (C_nar_) or widening (C_wide_) channels. (C) Five *Per* blocks were interspersed with four *Obs* blocks in a session.

**Figure 2.**
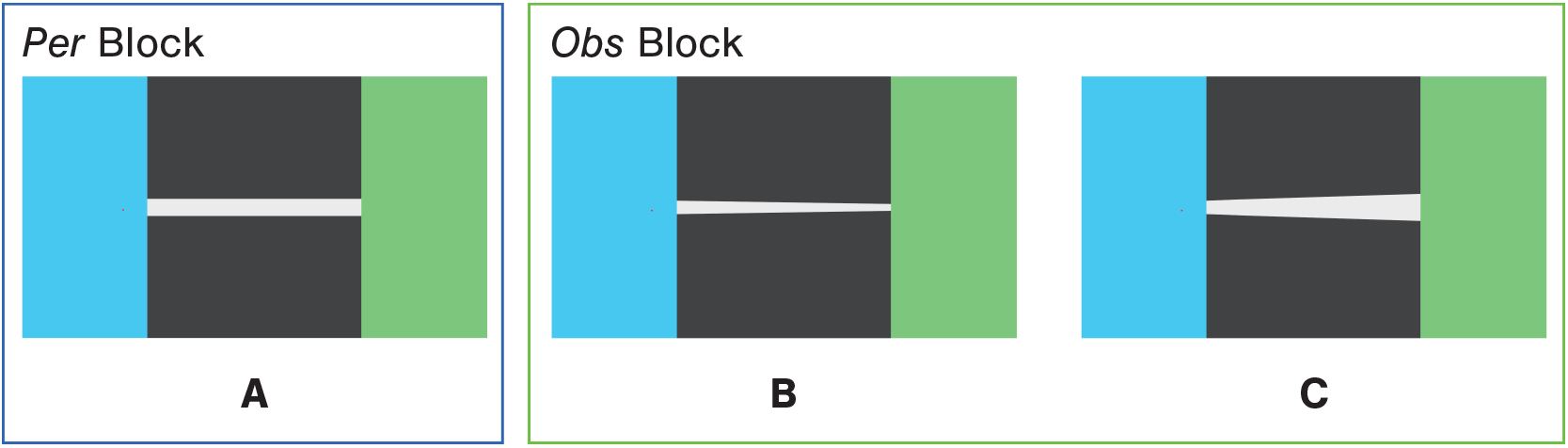
The appearance of the channels presented in the *Per* block (A: constant channel, C_const_) and the *Obs* block (B: narrowing channel, C_nar_; and C: widening channel, C_wide_).

**Figure 3.**
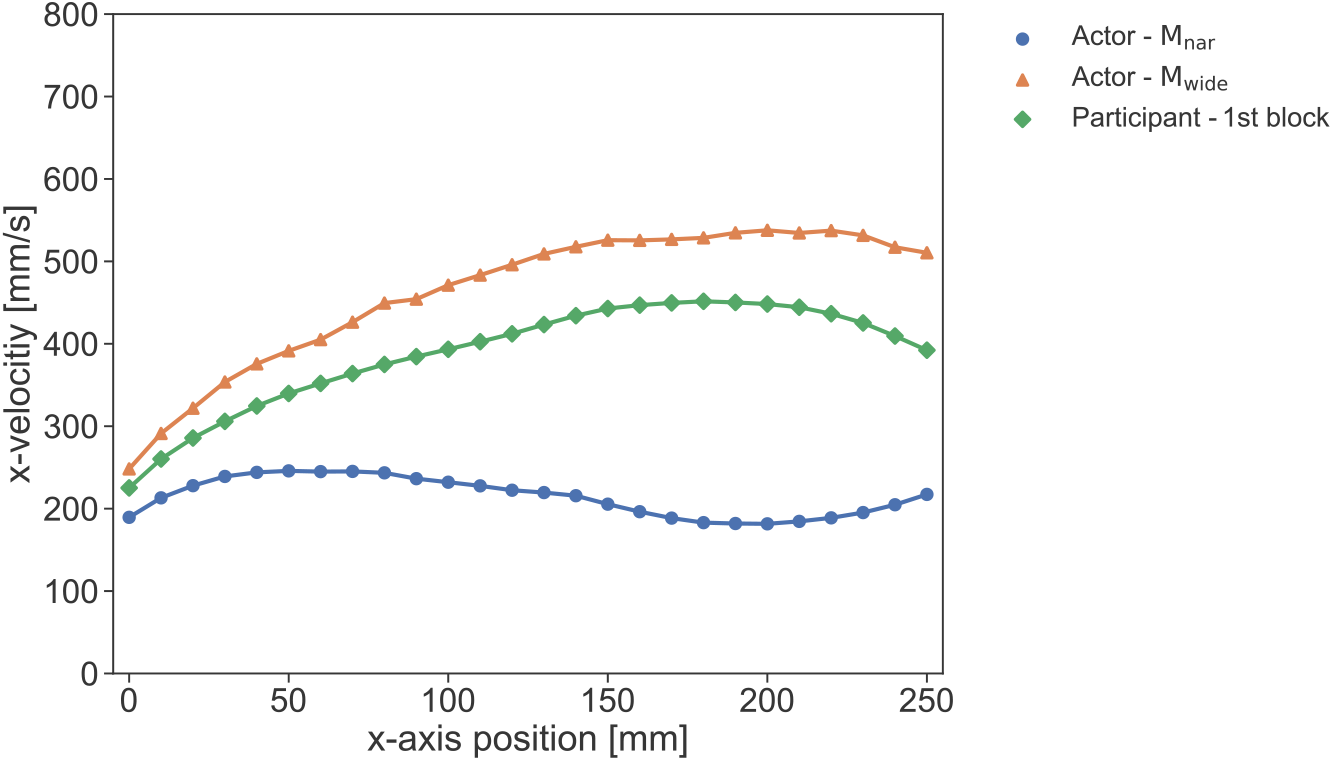
The x-velocity along the channel passage of the participants’ first *Per* block of the first session and the actors’ movement in each condition of *Obs* blocks (i.e., M_wide_ and M_nar_) in Experiment-1. The plot shows the average velocity for every 10 mm of position in the channels’ x-direction.

## Results

### Experiment-1

Twenty-four participants took part in Experiment-1. The participants performed five performing blocks (hereafter *Per* block; Figure 1 A) that were interspersed with four observation blocks (hereafter *Obs* block; Figure 1 B) per session (Figure 1 C). In the *Per* blocks, participants were required to move a cursor through a straight channel of a constant width (C_const_; see Figure 2 A) on a screen using a haptic stylus. The participants worked in three C_const_ different (constant) widths and were required to move through as quickly as possible without touching the walls. The participants had to make nine cursor movements (three times per channel in a randomized order) in each *Per* block. In the *Obs* blocks, the participants watched videos of the cursor movements by unknown actors.

The participants took part in four sessions with identical *Per* blocks. In each session, they watched one of the two movements (M_wide_ or M_nar_) in one of the two channels (C_wide_ or C_nar_, see Figure 2 B, C), as shown in Table.1. Each trial of the *Obs* block started with the presentation of a fixation point in the center of the monitor. The participants were asked to maintain their gaze on this point till the start of the recorded cursor movement video. The participants were required to keep their eyes on the cursor during the video presentation.

**Table 1.** Experiments-1 consists of four sessions of that combined the two channels (C_wide_ and C_nar_) and the two movement conditions (C_wide_ and C_nar_).

| Movements | Channels |  |
| --- | --- | --- |
| | Narrowing channel ($C_{nar}$) | Widening channel ($C_{wide}$) |
| Movement on $C_{nar}$ ($M_{nar}$) | $C_{nar}M_{nar}$ | $C_{wide}M_{nar}$ |
| Movement on $C_{wide}$ ($M_{wide}$) | $C_{nar}M_{wide}$ | $C_{wide}M_{wide}$ |

We analyzed the effects of the observation on participant’s own task performance in each of the *Per* blocks using a three-way (2 *channels* × 2 *movements* × 5 *blocks*) repeated measures analysis of variance (ANOVA). We expected to see the effects of AICs, and environmental effects (if any) as the main effects in the ANOVA, while we expected to observe PECs as an interaction of the channels and movements or as a three-way interaction in the ANOVA. The task performance of the participants was evaluated using the *error rate* (percentage of trials in which they touch the channel wall), the *peak distance* (maximum vertical distance of the pointer from the center of the channel), and the *movement time* or MT (time taken in each trial to pass through a channel).

First, we confirmed the change of velocity through the channel in each of M_wide_), M_nar_, and the participants’ initial *Per* block (i.e., the first block of the first session). M_wide_ tended to accelerate gradually during traversal through the channel, while Mnar did not show much velocity change but tended to decelerate slightly in the middle of the channel. Participants’ movements in the initial session also tended to accelerate gradually but not as much as M_wide_ (Figure 3). Participants’ movements in the initial session also tended to accelerate gradually but not as much as M_wide_. The average difference between the maximum velocity and the velocity at the start line was 310 (*s.d*. = ±53) mm/s for M_wide_, 79 (*s.d*. = ±27) mm/s for (M_nar_, and 257 (*s.d*. = ±133)mm/s for the participants’ first *Per* block (9 trials × 24 participants = 216 trials in total). In addition, the average MT was 560 (*s.d*. = ±51) ms for M_wide_, 1214 (*s.d*. = ±171) ms for (M_nar_, and 703 (*s.d*. = ±267) ms for the participants’ first *Per* block. Overall, Mwide had a higher velocity than the participants’ initial session, while Mnar had a slower velocity than the participants’ initial session.

Next, we analyzed the error rate. The participants made 4698 trials in total (including 378 failed trials). Because normality was rejected for some of the data groups, we conducted a three-way repeated measures ANOVA after carrying out the non-parametric aligned rank transform (ART) procedure (*α* = .05)^20^. The result did not show main effects for any of the factors (*channel*: *F*(1,23) = 0.637, *p* = .433, 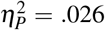; *movement*: *F*(1,23) = 0.155, *p* = .698, 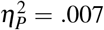; *block*: *F*(4,92) = 1.296, *p* = .277, 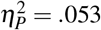). There were significant differences in the interaction between the *channel* and *block* and between the *movement* (*F*(4,92) = 2.961, *p* = .024, 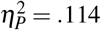) and *block* (*F*(4,92) = 3.294, *p* = .014, 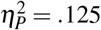), but not between the (*channel* and *movement*: *F*(1,23) = 0.320, *p* = .577, 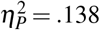; three-way: *F*(4,92) = 1.118, *p* = .353, 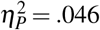). The results of the post-hoc analysis showed the simple main effect of the *block* only in the C_nar_ condition (*F*(4,92) = 3.074, *p* = .020, 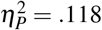), and the multiple comparisons (ART contrast test21) showed the significant difference between the third and fourth blocks (*t*(92) = −3.058, *adj.p* = .024). This may be due to the low error rate in the third block in the C_nar_ channel (see Figure 4 left). Overall, there was no significant difference in the error rates among the *channel* and *movement* conditions (see Figure 4 right).

**Figure 4.**
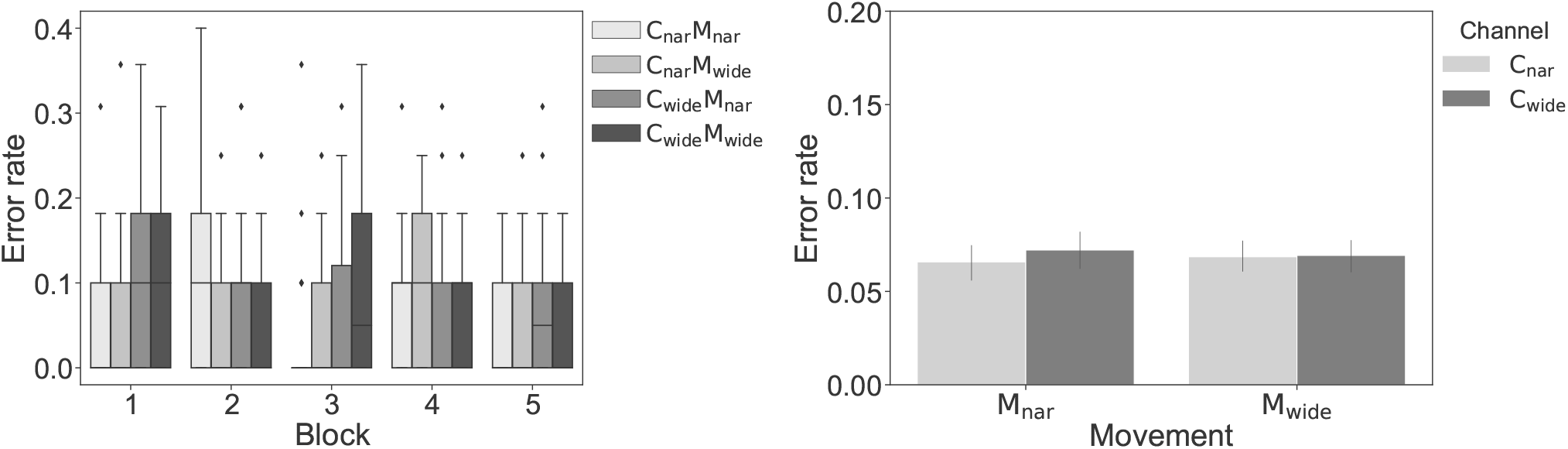
The error rate in Experiment-1. The left panel shows the error rate for each *Per* block in each condition. The right panel shows the average error rate across blocks 2 to 5 of each condition. Error bars show standard error.

Similarly, no significant difference was observed across the *channel* and *movement* conditions in the peak distance (Figure 5). Normality was not rejected under all the conditions (*p* > .05). The three-way ANOVA (*α* = .05) showed the significant main effect of only the *block* (*F*(4,92) = 2.647, *p* = .038, 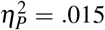). There were no other main effects (*channel*: *F*(1,23) = 0.145, *p* = .707, 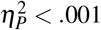; *movement*: *F*(1,23) = 0.435, *p* = .516, 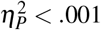) or interactions (*channel* and *movement*: *F*(1,23) = 3.134, *p* = .090, 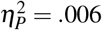; *channel* and *block*: *F*(4,92) = 0.615, *p* = .653, 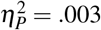; *movement* and *block*: *F*(4,92) = 0.109, *p* = .979, 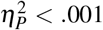; three-way: *F*(4,92) = 1.076, *p* = .373, 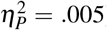). The results of the post-hoc analysis of the main effect of the *block* showed no significant differences between any of the blocks (*ad j.p* > .05). The p-value was adjusted using Holm’s sequential Bonferroni procedur^e22^. Overall, these results suggest that the observations in this experiment did not significantly affect the accuracy of the participants’ cursor manipulations.

**Figure 5.**
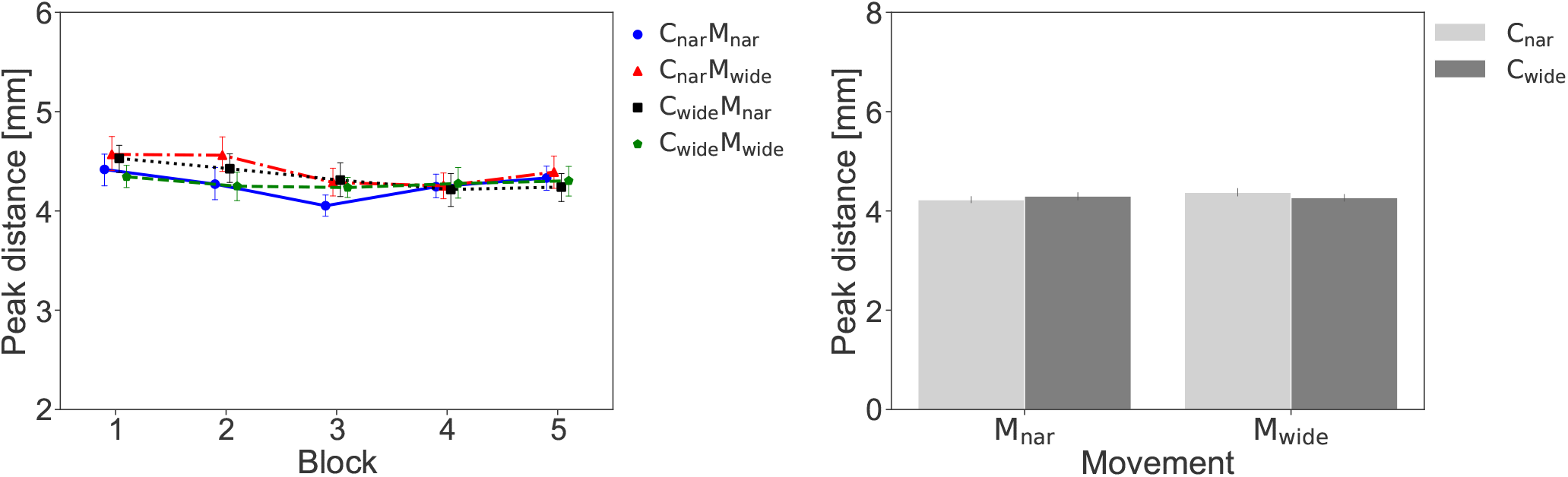
The peak distance from the center line of the channel during Experiment-1. The left panel shows the changes in the peak distance of the participants’ steering tasks across the *Per* blocks in each condition. The right panel shows the average peak distance across blocks 2 to 5 of each condition. Error bars show standard error.

However, the MT was found to be modulated by the *channel* and *movement* conditions (see Figure 6 left). Again, due to a lack of normality in some of the conditions, we conducted the ART ANOVA, which exhibited significant main effects for all the factors (*channel*: *F*(1,23) = 10.482, *p* = .004, 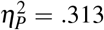; *movement*: *F*(1,23) = 18.664, *p* < .001, 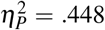; *block*: *F*(4,92) = 10.342, *p* < .001, 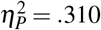) and for the interaction between the *movement* and *block* (*F*(4,92) = 6.114, *p* < .001, 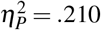). The interactions between the other factors were not significant (*channel* and *movement*: *F*(1,23) = 1.471, *p* = .238, 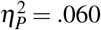; *channel* and *block*: *F*(4,92) = 0.875, *p* = .482, 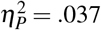; three-way: *F*(4,92) = 0.927, *p* = .452, 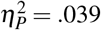). The results of the post-hoc analysis showed the simple main effect of the *block* only in the M_wide_ condition (*F* (4,92) = 12.130, *p* < .001, 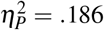) and the multiple comparisons (ART contrast test) showed the significant difference between the first and third blocks (*t*(92) = 6.020, *adj.p* < .001), first and fourth blocks (*t*(92) = 7.026, *adj.p* < .001), and first and fifth blocks (*t*(92) = 6.907, *adj.p* < .001). In addition, although the interaction between the *channel* and *movement* factors did not reach statistical significance in the three-way ANOVA, we performed multiple comparisons of the average MT in the four *Per* blocks after the *Obs* blocks (i.e., *Per* blocks 2 to 5) to observe the differences between the six pairs consisting of four combinations (Figure 6 right). The results showed the statistically significant differences in the average MT between C_wide_M_nar_-C_wide_M_wide_ (*adj.p* = .014, C_wide_M_nar_ > C_wide_M_wide_)’ C_wide_M_nar_-C_nar_M_wide_ (*adj.p* = .001, C_wide_M_nar_ > C_nar_M_wide_), C_wide_M_wide_-C_nar_ M_wide_ (*ad j. p* < .001, C_wide_M_wide_ > C_nar_M_wide_), and C_nar_M_nar_-C_nar_M_wide_ (*ad j*. *p* < .001, C_nar_M_nar_ > C_nar_M_wide_). This means that under the same *channel* conditions, the average MT is faster when observing Mwide than when observing M_nar_. In addition, it was significantly faster in the C_nar_M_wide_ condition than in any other condition.

**Figure 6.**
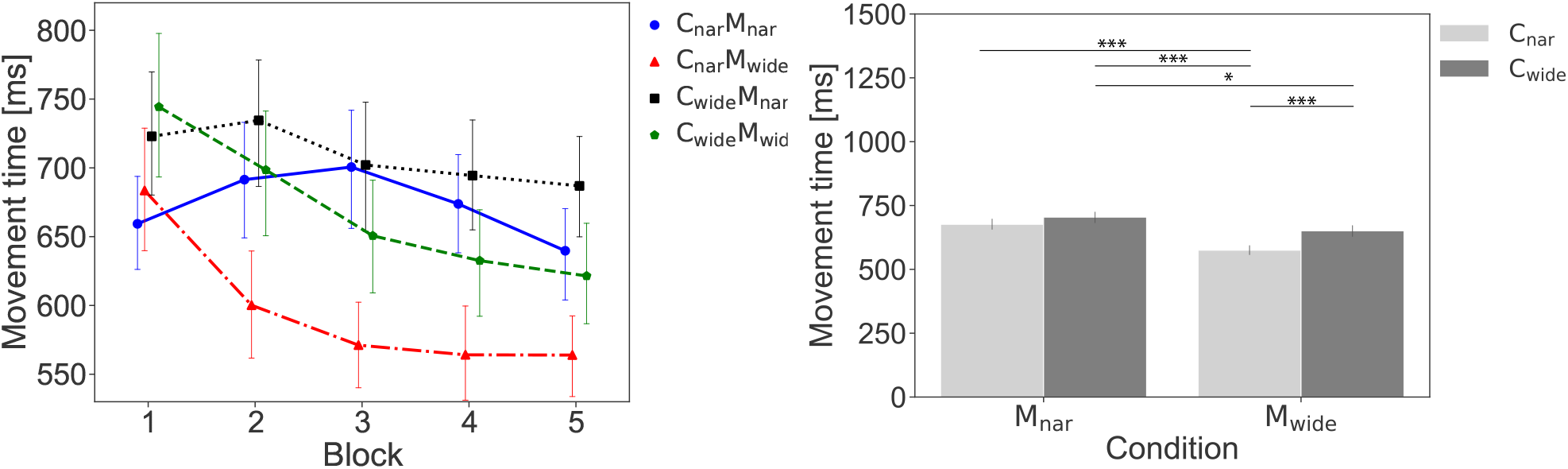
MT in Experiment-1. The left panel shows the changes in the participant MT across the *Per* blocks in each condition. The right panel shows the average MT across blocks 2 to 5 of each condition. Error bars show standard error. \**p* < .05,\*\**p* < .01, *p* < .001.

The above results suggest that while accuracy (i.e., error rate and peak distance) remain unaffected by observing actor movements and other channels (background environment), motor contagions are induced in one’s MT (speed) by observing others’ movements as well as environments. However, while we observed clear main effects of channel type and observed movement, we did not observe an interaction effect between the two. Thus, it remained unclear whether observing the channel itself affected the participants’ behavior or whether the cursor movement in the channel was required for the contagion to be induced. To confirm this, we conducted Experiment-2.

### Experiment-2

#### Twelve participants participated in Experiment-2

All of them had prior experience of Experiment-1. Procedures followed in Experiment-2 were the same as in Experiment-1, except for the fact that the participants observed only the C_wide_ and C_wide_ channels in each trial of the *Obs* blocks, and not the cursor movement within them. The participants attended just two sessions of the same *channel* conditions (i.e., C_nar_, C_wide_).

A total of 1151 trials were analyzed in Experiment-2 (including 71 failed trials). We confirmed that the *channel* conditions did not affect the error rate (Figure 7). The two-way ART ANOVA (2 channels × 5 blocks) showed the significant difference in the main effect of *block* (*F*(4,44) = 2.784, *p* = .038, 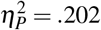). The main effect of channel (*F*(1,11) = 0.765, *p* = .400, 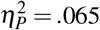) and interaction (*F*(4,44) = 0.284, *p* = .887, 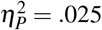) were not significantly different. Similarly, we did not observe any effect on the peak distance (Figure 8) in Experiment-2. The ART ANOVA showed no significant difference in any main effect or interaction (*channel*: *F*(1,11) = 0.005, *p* = .943, 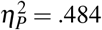; *block*:*F*(4,44) = 0.795, *p* = .535, 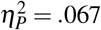; interaction: *F* (4,44) = 0.519, *p* = .722, 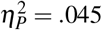).

**Figure 7.**
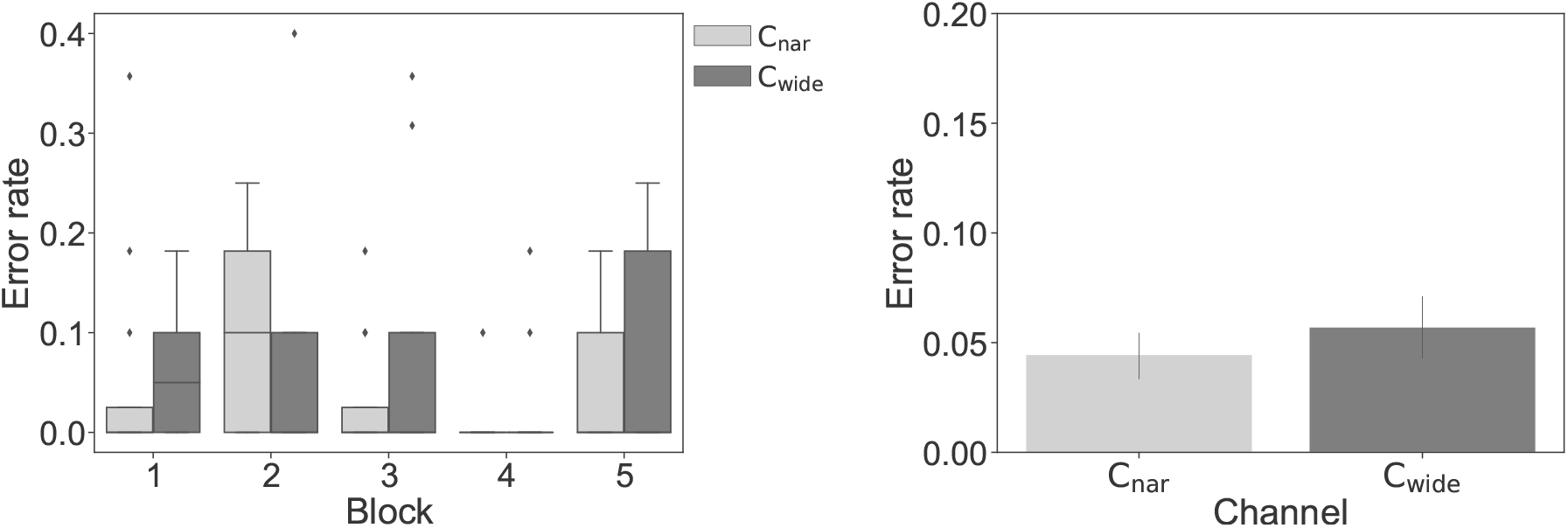
The error rate in Experiment-2. The left panel shows the error rate for each *Per* block in each condition. The right panel shows the average error rate across blocks 2 to 5 of each condition. Error bars show standard error.

**Figure 8.**
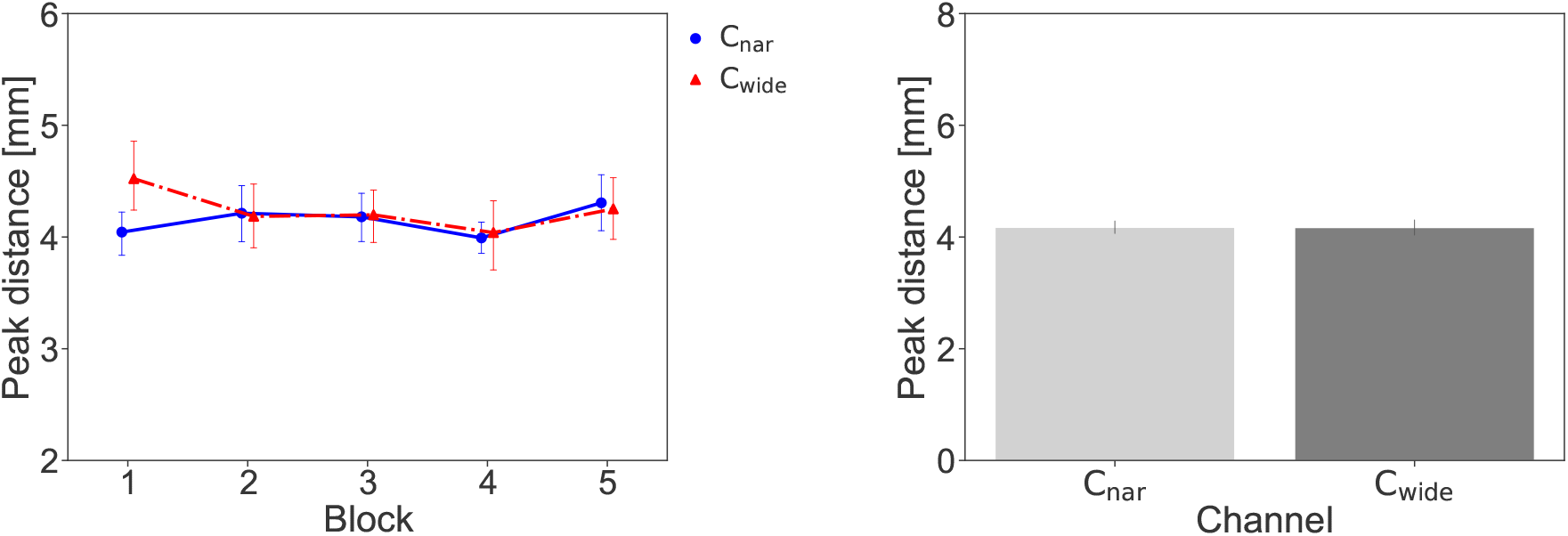
The peak distance from the center line of the channel during Experiment-2. The left panel shows the changes in the peak distance of the participants’ steering tasks across the *Per* blocks in each condition. The right panel shows the average peak distance across blocks 2 to 5 of each condition. Error bars show standard error.

In addition, we found no difference in the MT in Experiment-2 (Figure 9). The ART ANOVA showed no significant difference in any main effect or interaction (channel: *F*(1,11) = 0.058, *p* = .815, 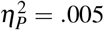; *block*:*F*(4,44) = 1.131, *p* = .355, 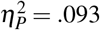; interaction: *F*(4,44) = 0.519, *p* = .722, 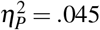). This result suggests that the difference in the MT between the channel conditions observed in Experiment-1 does not occur when the cursor movement through the channel is absent.

**Figure 9.**
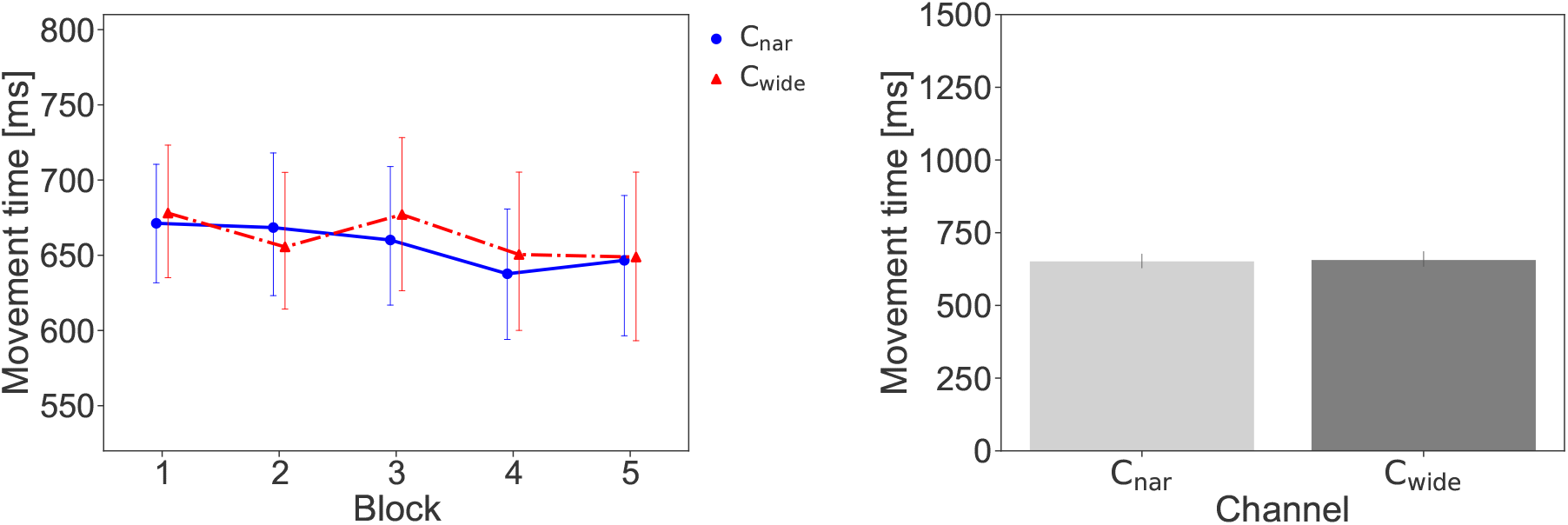
MT in Experiment-2. The left panel shows the changes in the MT of the participants’ steering tasks across the *Per* blocks in each condition. The right panel shows the average MT in blocks 2 to 5 of each condition. Error bars show standard error.

## Discussion

In two experiments, we explored whether and how the environment in the background of an action modulates the motor contagions in an observer. In Experiment-1, we observed that the MTs of the observers were influenced both by the movement of the cursor they observed, as well as by the background (channel shape) in which the cursor movement was observed. In Experiment-2 we showed that while the background influences the motor contagions, it does so only in the presence of the action in the foreground.

There are several possibilities why the background may have caused the differences in motor contagions. First, the channel shapes may have created a visual illusion and modified the perception of cursor speed within the channels. Specifically, the C_nar_ channel may have led to a general increase in the perceived speed of the cursor, leading to an increase in the observer’s own steering speed after observing movement in the C_nar_ (Figure 6). Similarly, the C_wide_ channel may have led to an overall decrease in the perceived cursor speed, leading to a decrease in the observer’s own steering speed after observing the movement in the Cwide. Second, the background may have modulated the competitive motivation of observers, making them copy the performance by the actors. That is, observing an accelerated movement (M_wide_), which is typical of the C_wide_ channel, in the C_nar_ channel could have motivated the participants to quicken their own movements. Conversely, observing a deaccelerated movement (Mnar) in the C_wide_ channel could have induced the participants to slow their own movements. This possibility is supported by our measure of the index of performance, which is a generalized performance index independent of the path conditions^23–26^. We observed a strong correlation between the index of performance of the observed and performed actions by the participants (see Figure S1 in the Supplementary Information for more details). However, given that few data points were available for this analysis, further studies are required to confirm performance copying as the cause of the results observed.

We note here that we did not observe any PECs in our study. Apriori, according to the findings of previous studies^7, 8, 12^ we expected PECs to manifest as acceleration changes opposite to the observations. That is, we expected the participants to decrease their own movement speed when they observe an accelerated movement in the C_nar_ channel (when in fact one would expect a deaccelerated movement in it) and conversely increase their movement speed after watching deaccelerated movements in the C_wide_ channels. Differences in the experimental design may explain the lack of PECs in this study. Specifically, PECs are known to be heavily modulated by the perceived goal of the actor, and the participants in this study were not instructed in detail about the intention or goal of the actor because here we focused on the environmental effects rather than the task instructions. In addition, the participants’ own movements were presented on screen in real-time, and the presence of visual feedback in addition to the AIC an environment related contagions may have hidden any PECs induced in the participants’ behaviors.

However, prediction errors did modulate our results. In this paper, we focused primarily on environmental effects, and hence we considered a three-way ANOVA of MT across movement types, channel types, and blocks (Figure 6). On the other hand, the MT may be instead analyzed in the presence of prediction error (when observed movement and channel do not match) and absence of prediction error (when observed movement and channel match) (see Figure S2 in the Supplementary Information). We observed a significant interaction of MT between movement type and prediction error (*F*(1,23) = 9.507, *p* = .005, 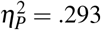) and that in the presence of prediction error, the AIC effect tends to increase. Several previous studies have reported that unexpected visual stimuli increase human attention^27, 28^ and we believe attention difference was also the reason for the prediction error related increase observed in our study. However, further studies are required to verify this issue.

In our experiments, the MT changes were sustained despite the visual feedback of the cursor. However, it remains unclear if the participants perceive this MT change in their movement. The answer to this question is interesting in regard to a sense of agency (or the subjective perception of controlling one’s action) perceived by the participants towards their movements^29–33^ when suffering from motor contagions. Several previous studies suggest the sense of agency to be modulated by prediction errors generated by the internal models used for our movements^34, 35^, and evaluating the effects of motor contagions on the sense of agency may be a way to quantify the effect induced by motor contagions on the learned internal models driving one’s movements.

A previous study that reported both AICs and PECs observed that motor contagion occurred even when the observing agent was instructed to aim for a goal different from their ow^n8^. However, the tendency of behavior change varied depending on whether the observed goal was consistent with the observer’s own goal or not^8^. This suggests that the environmental effects may also differ depending on how similar the observed environment is to one’s own task environment. In our current study, we did not modulate the environment enough to answer this question, but hope to clarify the influence of the relationship between the observed and the observer’s environment in future studies.

The finding that the environment can affect motor contagions is novel and important in the design of systems that support training through observational motor learning^36, 37^ and driving users’ movements implicitly by using apparent motion^38^. A unique feature of our experimental results is that the CnarMwide condition modulated the movements in a direction that improved task performance (like Vasalya et al.39). No effect of the observation on the error rate or peak distance was observed and the MTs shortened with repeated observations. The average MT was significantly shorter than that of the C_wide_M_wide_ condition, in which the observed motion itself was the same. These results show that motor contagions can be used to improve individual motor performance and that the environment is one of the critical factors in observational motor learning.

## Methods

### Task and apparatus

Our participants were required to pass the red cursor (2 mm in diameter) through the light gray channel and then move it from the blue area on the left to the green area on the right. The length of the channel was fixed at 250 mm throughout the experiment. The width of the channel changed in each trial (see the next subsection for more details). The participants were required to pass the cursor as quickly as possible without exceeding the upper and lower boundaries.

The participants sat facing an LCD monitor (Dell, 240 Hz, 24.5 inches) that presented visual information. They held the stylus of a 3D haptic device (3D Systems Touch) with their dominant hand. Programmed forces restricted the haptic stylus movement in a plane parallel to the computer screen such that the participants could manipulate the cursor on the screen by moving the stylus in a plane parallel to the computer screen. The cursor movement on the screen was scaled to match the physical movement of the stylus. The participants’ movements were recorded by the pointing device’s encoder at a sampling rate of 1000 Hz. The latency of the real-time visual feedback was 17 – 21 ms based on a 240 fps simultaneous video recording of the hand motion and visual feedback on the screen using an external camera.

### Participants

#### Experiment-1

Twenty-four naïve participants (17 males and 7 females; aged 19 - 36, mean age 24.4 years, standard deviation 4.19 years) took part in Experiment-1. This sample size was based on that calculated using G * Power 3.1 with a repeated measures ANOVA within factors, *α* = 0.05, *β* = 0.80, *f* = 0.25 (medium value), number of measurements = 4, correlation among repeated measures = 0.5, non-sphericity correction = 1^40^. All the participants had normal or corrected-to-normal vision and no disability. All the experiments were approved by the Life Science Research Ethics and Safety Office at the University of Tokyo, Japan (approval number: 19-414). All the experimental procedures described below were conducted in accordance with the guidelines of the Declaration of Helsinki41 and the procedure approved by the above ethics committee. Participants signed an informed consent form before taking part in the experiment and received a compensation of JPY 2,100 after completion.

#### Experiment-2

Twelve participants (9 males and 3 females; aged 20-31, mean age 24.3 years, standard deviation 4.03 years) took part in the Experiment-1. All of them had previous experience of Experiment-1. Experiment-2 was conducted on the same day as Experiment-1.

#### Experimental design in Experiment-1

In an experimental session, five *Per* blocks (Figure 1 A) were interspersed alternately with four *Obs* blocks (Figure 1 B), as shown in Figure 1 C. In the *Per* block, the participants performed the steering task themselves; in the *Obs* blocks, they observed the recorded images of the steering task being performed by unknown actors. We investigated the effect of the interaction between the observation of the motion and the observation of the environment (i.e., the channel shape) on the participant’s performance. Each participant took part in a session of the experiment in each of these conditions.

The conditions of the observation were set by combining the *channel* (C_nar_, C_wide_) and the *movement* of the cursor (M_nar_, Mwide) presented at *Obs* blocks of each session.

In the *Obs* blocks, the following two types of *channel* conditions for the observation, where the channel width changes linearly from the start line (i.e., the boundary with the blue area on the left) to the endpoint (i.e., the boundary with the green area on the right), were adopted:

- C_nar_: The channel width at the start line was 16 mm and the width at the end line was 8 mm (i.e., the channel width at the end line was half of the width at the start line; Figure 2 B).
- C_wide_: The channel width of the start line was 16 mm and the width of the end line was 32 mm (i.e., the channel width at the end line was twice the width at the start line; Figure 2 C).

As the *movement* conditions, the following two types of recorded data of the actor’s cursor motion were adopted:

- M_nar_: The recorded data when the actor passed the cursor through the C_nar_.
- M_wide_: The recorded data when the actor passed the cursor through the C_wide_.

The conditions of the observation were a combination of the two *movement* × two *channel* conditions. In other words, in the C_nar_M_nar_ and C_wide_M_wide_ conditions, the cursor movements were presented in the same channel as used to record the actor. On the contrary, in the C_wide_M_nar_ and C_nar_M_wide_ conditions, the cursor movements were presented in a different channel from that used to record the actor.

In the *Per* blocks, the channel width was constant from the start to the endpoint, as shown in Figure 2 A. The channel width was selected from the following three conditions by a pseudo-random algorithm in each trial: 12 mm, 16 mm, and 20 mm.

### Procedure in Experiment-1

Before the main sessions, the participants took part in training sessions. After that, they participated in four sessions for each of the above mentioned conditions. The order of sessions was counterbalanced between the participants. Each session involved five *Per* blocks and four *Obs* blocks. The participants had a break of three minutes at least between the sessions.

#### Training sessions

The participants took part in three types of training sessions to practice the steering task. In the first training session, they started with a wide C_const_ (24 mm), and as they became used to the operation, they experienced tasks with narrower C_const_ (12 mm at the end of the session). In this session, the participants were able to confirm the operation and their comfortable posture to become sufficiently familiar with the task. In the second training session, the participants experienced tasks with C_wide_ and C_nar_. We conducted this session to give participants an experiential understanding of the difficulty of the tasks for each channel for the observation. In the final training session, the participants experienced the tasks in the three conditions of C_const_ in an order determined by a pseudo-random number, as in the *Per* blocks. This session included nine trials (three trials for each C_const_).

#### Per blocks

In the *Per* blocks, all the participants were required to perform the steering task as quickly as possible, and with as few failures as possible. Before the trial, the C_const_ of this trial and cursor were displayed. Further, the stylus was fixed to the start position in the blue area. Then, the participant started the trial by pressing the button on the stylus with their thumb. During the trial, the stylus could be moved freely up, down, left, and right. The participant manipulated the stylus to make the cursor pass through the channel by holding down the button. At this time, a clamping force was presented in the depth direction to guide the manipulation in a plane parallel to the monitor. When the cursor reached the green area, a sound was presented to indicate success. Then, when the participant released the button, the stylus was locked in its current position. If the participant touched the upper or lower border while passing through the channel, a sound indicating failure was presented, and the stylus was fixed in that position. Then, the display blacked out and the stylus returned to the initial position automatically. When the stylus reached the initial position, it was fixed and the channel of the next trial and cursor were displayed.

The participants performed nine trials of the *steering task* in each block. If the participant touched the upper or lower boundary while passing through the channel, it was judged as a failed trial and was not counted. A trial with the same channel width condition was added to the end of the block. In other words, the participant had to succeed in the steering task nine times (i.e., three times for each channel width) per block.

#### Obs blocks

In *Obs* blocks, the participants were required to observe the cursor motion performed by the actors. In each *Obs* block, nine samples were randomly selected out of 30 samples (3 actors × 10 samples) of motion data for the *movement* condition in that session (i.e., M_wide_ or M_nar_). Before the start of the first *Obs* block, the participants were informed of the channel conditions that would be presented in the session (i.e., C_wide_ or C_nar_). They were not informed about the *movement* condition. When the trial started, a black background and a cross-shaped gazing point at the center of the screen were shown for 1000 ms. After that, the gazing point disappeared, a channel and a cursor appeared, and the cursor moved from left to right. After the cursor reached the area on the right, the screen blacked out and the next trial started after a 1000 ms blank. The participants were required to keep their eyes on the gazing point at the start of the trial and on the cursors passing through the channel during the trial.

#### Obs block videos

To create the video image for the *Obs* blocks, the two-dimensional coordinates of the cursor, manipulated by the actor, were recorded. The three actors were given the same basic task instructions as the participants and repeatedly performed steering tasks in the C_wide_ and C_nar_ conditions. The coordinate data were recorded at a sampling frequency of 240 Hz in each trial using the same equipment as in the experiments. The recording for each actor continued in the trials until 10 samples per channel condition were collected. We adopted only those motion data that fell within the Cnar) condition for the Mwide condition. This was because the main experiment had a condition in which the Mwide condition was presented in the Cnar) condition (i.e., CnarMwide). In other words, for the trials in the C_wide_ condition, even if they were successful, the data were not collected if they did not fall within the C_nar_ condition. However, during the recording, whether the data had been collected was not fed back to the actors. Therefore, the motion speed was not intentionally suppressed to keep the actor within the C_nar_ condition.

### Experimental design and procedure in Experiment-2

In Experiment-2, the participants observed only the channel without the moving cursor in the *Obs* blocks. The two *channel* conditions were the same as in Experiment-1 (i.e., C_nar_, C_wide_). The participants took part in two sessions for each of the conditions. The session consisted of five *Per* blocks and four *Obs* blocks, as in Experiment-1. The order of the sessions was counterbalanced between the participants.

In each *Obs* block, the following trial was conducted nine times:

- A black background with a cross-shaped gazing point placed at the center of the screen was first presented for 1000 ms.
- Then, the gazing point disappeared and the image of the channel (C_nar_ or C_wide_) was presented for 1500 ms.
- The screen was finally blacked out, and a 1000 ms blank screen was presented until the next trial.

The participants were required to keep their eyes on the gazing point at the start of the trial and on the channel during the trial. All the other task designs and procedures were the same as in Experiment-1.

### Data analysis

The error rate and maximum vertical peak distance were measured as indicators of task accuracy. The error rate is the number of failed trials in blocks 2-5 of each session divided by the total number of trials in blocks 2-5. The peak distance was the maximum vertical offset of the cursor trajectory from the center of the channel during each trial. Only successful trials were included in the peak distance analysis; unsuccessful trials were excluded. The cursor trajectory from the start line to the end line was recorded using the input device’s encoder (position resolution 0.055 mm) at a refresh rate of 1000 Hz. Of the recorded samples of the cursor position for each trial, the farthest vertical distance from the center line was defined as the peak distance. The time from passing the start line to passing the end line in a successful trial was measured as the MT.

Statistical analyses were performed for the three parameters listed above (i.e., error rate, peak distance, and MT). In Experiment-1, we analyzed the effects of the observation on a participant’s task performance in each of the*Per* blocks using three-way (2 channels × 2 movements × 5 blocks) repeated measures ANOVA. In Experiment-2, we conducted two-way (2 channels × 5 blocks) repeated measures ANOVA. Before the ANOVA, we confirmed the normality of the data using the Shapiro-Wilk test. Then, if normality was rejected, we conducted the ANOVA using the ART procedure^20, 42^. The post-hoc analysis was conducted using pairwise t-test for the normal data, and the ART procedure for multifactor contrast tests21 for the non-normal data. The p-value was adjusted using Holm’s sequential Bonferroni procedur^e22^. In addition, to examine the differences between each condition of the four combinations (i.e., C_wide_M_nar_, C_wide_M_wide_, C_nar_M_nar_, and C_nar_M_wide_), multiple pairwise Wilcoxon signed rank exact test (using Holm’s sequential Bonferroni procedure) wad conducted on the average values of *Per* blocks 2 - 5 for the MT in Experiment-1.

### Steering task

To examine the relationship between the environment and movement in the motor contagions, we employed a behavioral task called the *steering task* proposed by Accot and Zhai^18, 19, 24^. In this task, users manipulate a visual object such as a mouse cursor to pass through a narrow path. In this steering task, the so called *steering law model*^18, 19, 24^, defines the user performance. This *steering law model* is derived from *Fitts’ law*^43^ and is confirmed to fit various conditions and GUI operations such as indirect-input stylus^18^, mouse, touchpad, trackball, track point^19^, direct-input pen tablet^44^, and 3D pointing device with haptic feedback^45^. According to this model, the relationship between the MT required to traverse straight channel with a specific amplitude (*A*) and width (*W*) is represented by the following expression (Figure 10 upper image)^18^:

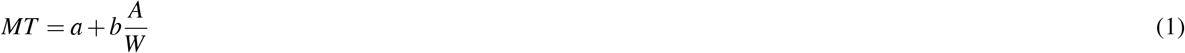

*a* and *b* are empirically determined constants, while *A/W* is also known as the index of difficulty (*ID*). Hence, as the amplitude lengthens or the width narrows, the difficulty of passing through the channel increases, and the MT lengthens. The average cursor speed (*V*) is represented by the following expression^18^:

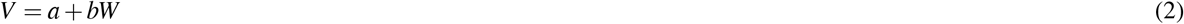

This indicates that the speed of passing through the channel depends on the channel width and that it quickens as the width widens and slows as the width narrows.

**Figure 10.**
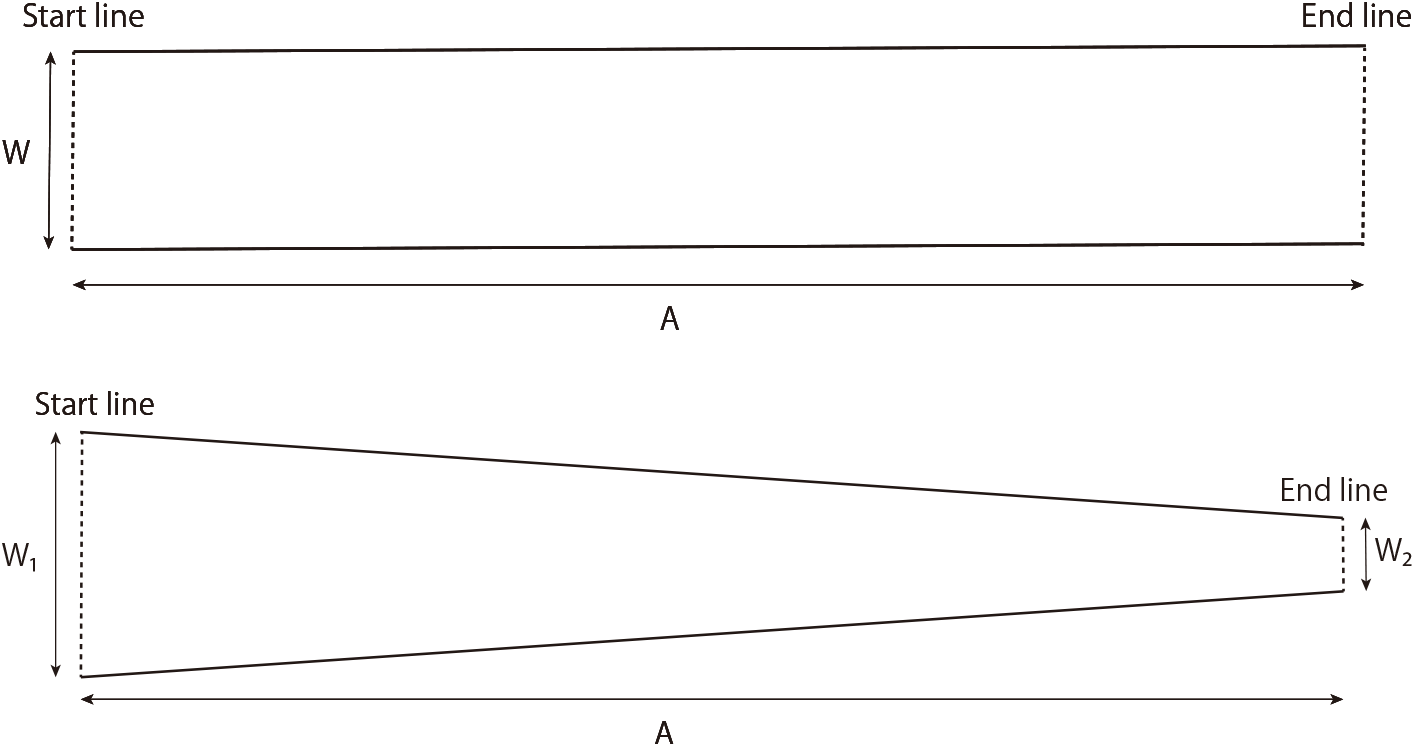
The straight channel (upper image) and narrowing channel (lower image) used in the steering task^18^

It has been suggested that the *ID* can be defined not only for the straight channel with a constant channel width, but also for various channel conditions. It has also been suggested that the MT can be predicted by a model based on the *ID*, as shown in the following expression^18^:

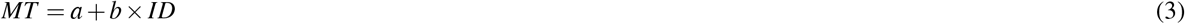

For example, for a gradually narrowing channel, the *ID* can be defined using the channel widths at the start and end lines of the channel (see *W*_1_ and *W*_2_ of Figure 10 lower image), as in the following expression^18^:

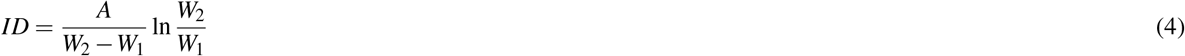

In addition, Yamanaka et al. conducted an experiment that included a gradually widening channel with *W*_1_ and *W*_2_ inverted above the narrowing channe^l44^. Their experimental results suggest that the speed of passing cursors is progressively slower in narrowing channels and progressively faster in widening channels, although corrections to the model depending on the orientation of the channel may allow for more accurate predictions of the MT.

## Acknowledgments

This research was supported by JSPS KAKENHI Grant Number 21K17786, JST ERATO Grant Number JPMJER1701, and JST Moonshot R&D Program Grant Number JPMJMS2292, Japan.

## Author contributions statement

H.S. K.F., and G.G. conceived and designed the experiments. H.S. conducted the experiments. H.S. and G.G. analyzed the data. H.S. and G.G. contributed to the preparation of the manuscript. H.S. K.F., M.I., and G.G. reviewed the manuscript.

