## Supplementary Information for "Background environment modulates motor contagions in humans"

### Supplementary figure

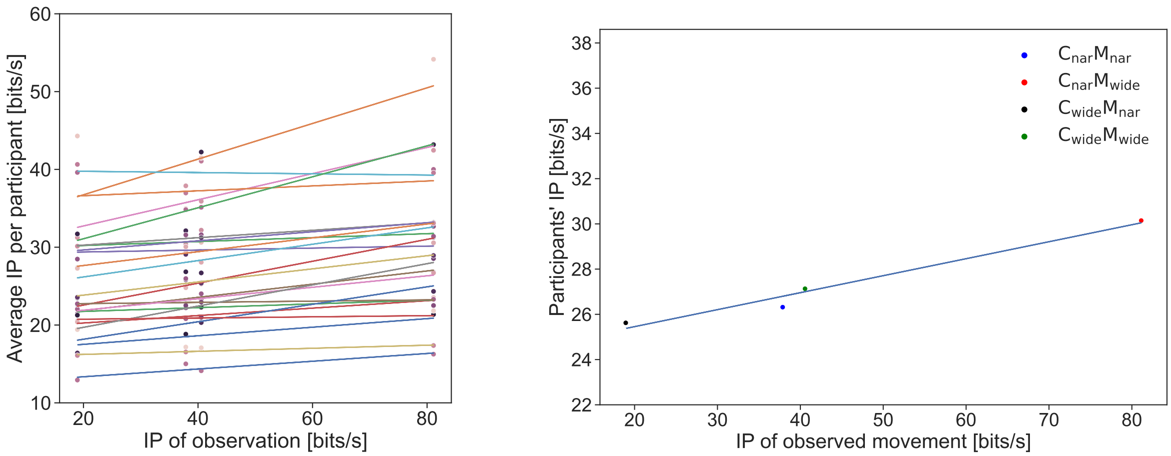


**Figure S1.** The relationship between the index of performance (*IP*) of the observed movements and *IP* of the participant's movements in Experiment-1. The *IP* is a generalized performance index independent of the path conditions in *Fitts' law* and the *steering law*, as shown in the following expression:

$$IP = ID/MT$$

. The graph is a scatterplot with the x-axis as the IP of the observed movement in the *Obs* block and the y-axis as the IP of the participant in the *Per* block after the observation (i.e., blocks 2–5; left: mean value of each participant, right: mean value of all the participants). The straight line represents the regression line. Although the number of points is insufficient to guarantee the results' reliability, the plot of the mean value of all the participants (i.e., right graph) shows a positive correlation between the participants' IP and the IP of the observed movements ($y = 0.075x + 23.954$, $R^2 = 0.973$).

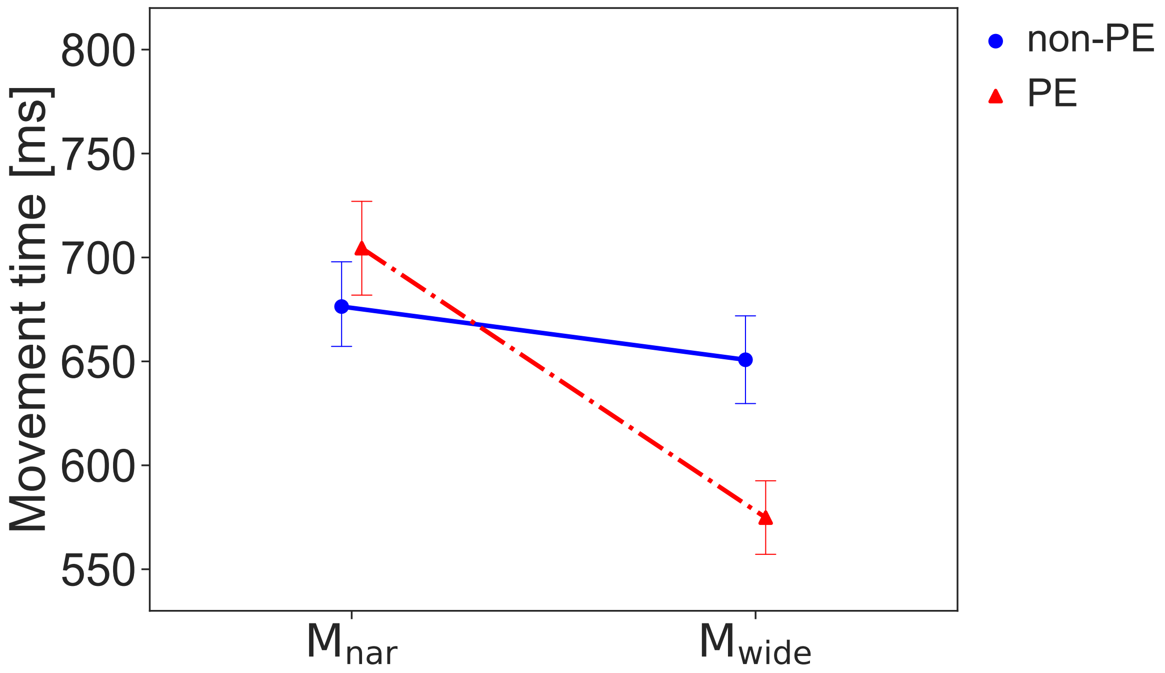


**Figure S2.** The figure shows the MT data from Figure 6 replotted across movement type and prediction error (instead of channel type). We expected participants to have experienced a prediction error (PE) when the channel type did not match the movement type they observed. And we expected participants to not have experienced a prediction error when the channel type matched the movement type they observed (non-PE). The plot shows the average MT in blocks 2 to 5 of each condition in Experiment-1. Error bars show standard error. Normality was not rejected, and the two-way repeated measures ANOVA (2 *predictability* $\times$ 2 *movements*) showed a significant main effect of *movement* ($F(1,23) = 33.581$, $p < .001$, $\eta_P^2 = .594$) and interaction ($F(1,23) = 9.507$, $p = .005$, $\eta_P^2 = .293$). The main effect of *predictability* was not significant ($F(1,23) = 3.434$, $p = .078$, $\eta_P^2 = .129$).
